# Single cell analysis of the effects of developmental lead (Pb) exposure on the hippocampus

**DOI:** 10.1101/860403

**Authors:** Kelly M. Bakulski, John F. Dou, Robert C. Thompson, Christopher Lee, Lauren Y. Middleton, Bambarendage P. U. Perera, Sean P. Ferris, Tamara R. Jones, Kari Neier, Xiang Zhou, Maureen A. Sartor, Saher S. Hammoud, Dana C. Dolinoy, Justin A. Colacino

## Abstract

**Background:** Lead (Pb) exposure is ubiquitous and has permanent developmental effects on childhood intelligence and behavior and adulthood risk of dementia. The hippocampus is a key brain region involved in learning and memory, and its cellular composition is highly heterogeneous. Pb acts on the hippocampus by altering gene expression, but the cell type-specific responses are unknown.

**Objective:** Examine the effects of perinatal Pb treatment on adult hippocampus gene expression, at the level of individual cells, in mice.

**Methods:** In mice perinatally exposed to control water (n=4) or a human physiologically-relevant level (32 ppm in maternal drinking water) of Pb (n=4), two weeks prior to mating through weaning, we tested for gene expression and cellular differences in the hippocampus at 5-months of age. Analysis was performed using single cell RNA-sequencing of 5,258 cells from the hippocampus by 10x Genomics Chromium to 1) test for gene expression differences averaged across all cells by treatment; 2) compare cell cluster composition by treatment; and 3) test for gene expression and pathway differences within cell clusters by treatment.

**Results:** Gene expression patterns revealed 12 cell clusters in the hippocampus, mapping to major expected cell types (e.g. microglia, astrocytes, neurons, oligodendrocytes). Perinatal Pb treatment was associated with 12.4% more oligodendrocytes (*P*=4.4×10^−21^) in adult mice. Across all cells, differential gene expression analysis by Pb treatment revealed cluster marker genes. Within cell clusters, differential gene expression with Pb treatment (q<0.05) was observed in endothelial, microglial, pericyte, and astrocyte cells. Pathways up-regulated with Pb treatment were protein folding in microglia (*P*=3.4×10^−9^) and stress response in oligodendrocytes (*P*=3.2×10^−5^).

**Conclusion:** Bulk tissue analysis may be confounded by changes in cell type composition and may obscure effects within vulnerable cell types. This study serves as a biological reference for future single cell studies of toxicant or neuronal complications, to ultimately characterize the molecular basis by which Pb influences cognition and behavior.

## Introduction

Lead (Pb) exposure remains ubiquitous in many municipalities and rural areas. The removal of Pb from paint and gasoline has been a major public health success (Needleman et al. 1990), though Pb’s persistence in soil, dust, and historic house paint make abatement from our lives and environments difficult (Dissanayake and Erickson 2012). The Centers for Disease Control and Prevention estimate that approximately 500,000 children ages 1-5 in the United States have blood Pb levels above the reference level (≥ 5ug/dL) (Raymond and Brown 2017), and no safe level of Pb has been identified. Developmental Pb exposure has profound, permanent effects on behavior and cognition (Bellinger et al. 1984; Needleman et al. 1990; Sanders et al. 2009). Young children are particularly susceptible to Pb’s effects due to 1) 4–5 times higher Pb absorption than non-pregnant adults (WHO 2010); 2) higher Pb intake per unit body weight; 3) incomplete blood–brain barrier development and 4) neurological effects which occur at all levels of exposure (WHO 2007). Early life Pb exposure results in permanent cognitive and behavioral changes.

Many brain regions are influenced by Pb exposure (Sanders et al. 2009). In particular, the hippocampus, a brain region in the temporal lobe that is central to long-term memory formation and an important part of the limbic system (responsible for emotional regulation), is a key target of Pb’s effects. Magnetic resonance spectroscopy of the hippocampus of men with high cumulative Pb exposure revealed higher myoinositol-to-creatinine ratios with exposure (Weisskopf et al. 2007), suggesting that the hippocampus may have cellular changes in response to Pb. Toxicology studies have shown RNA expression differences in the bulk hippocampus with both acute and chronic Pb exposures (An et al. 2014; Schneider et al. 2012). For example, in mice, three days of intraperitoneal Pb exposure (15 mg/kg body weight) starting at postnatal day 12 was associated with hippocampal microglial activation and inflammatory cytokine generation (Liu et al. 2015). Multiple brain regions are influenced by Pb exposures, and the hippocampus has emerged as a brain region. where consistent Pb-driven changes were observed.

Bulk brain region studies are important first steps to characterize relevant exposure levels and disease endpoints (Bakulski et al. 2012; Malloy et al. 2019). The next challenge in developmental toxicology studies is distinguishing whether the persistent molecular effects identified are due to global changes, altered cellular proportions, or cell type specific changes (Jaffe and Irizarry 2014). The hippocampus is a highly heterogeneous structure, containing many cell types such as neurons, astrocytes, oligodendrocytes, and microglial cells. Recently, single cell RNA profiling through droplet-based techniques has allowed for the deep unbiased profiling of thousands of cells in the hippocampus. Use of this technique showed that the hippocampus has even more cellular heterogeneity than was initially thought (Stahl et al. 2016). As all of the previous in-depth molecular profiling of Pb exposed brain tissues utilized bulk tissues analyses, the majority of our knowledge comes from averaging across this complex heterogeneity. Identifying cell type-specific effects of developmental Pb exposure has potential long-term clinical benefits.

Understanding the cell type specific impacts of Pb exposure is critical to elucidate mechanisms of its effects and reveal possible avenues of prevention and treatment of adverse outcomes due to Pb exposure. Here, we apply single cell RNA profiling to comprehensively characterize the effects of perinatal Pb exposure on the adult hippocampus in a cell-specific manner in mice. Our results reveal global RNA changes after Pb exposure, consistent with prior research. We also identify oligodendrocytes and microglia as particularly vulnerable cell types to Pb exposure, demonstrating cell type-specific RNA changes and an exposure-dependent shift in cell type representation.

## Methods

### Animal Exposure Paradigm

The mice for this experiment derive from a colony maintained for over 230 generations where the *A^vy^* allele is passed through the male line. This results in forced heterozygosity on an invariant genetic background which is approximately 93% identical to the C57BL/6J strain (Waterland and Jirtle 2003; Weinhouse et al. 2014). Post-pubertal virgin *a*/*a* females (~5 weeks old) were randomized into exposure groups: control or drinking water containing 32ppm Pb-acetate. This dose and route of exposure was designed to be relevant to human perinatal exposure. Pb-acetate water was made following our previously described protocol (Faulk et al. 2014). Throughout the experiment, mice were maintained on a phytoestrogen-free modified AIN-93G diet (TD.95092, 7% Corn Oil Diet, Harlan Teklad).

After an initial two weeks of exposure, females were mated with *a*/*a* males. Exposure through drinking water continued through gestation and lactation. After weaning, 1 male and 1 female wildtype (*a*/*a*) offspring per litter were maintained on Pb-free drinking water. At 5 months, mice were sacrificed for experimental analyses. All animals had access to food and water *ad libitum* throughout the experiment, were housed in polycarbonate-free cages, and were maintained in accordance with the guidelines set by the Institute of Laboratory Animal Resources. The study protocol was approved by the University of Michigan Institutional Animal Care and Use Committee.

### Hippocampus Isolation and Dissociation

Immediately following euthanasia with CO_2_, mice underwent whole-body perfusion with cell-culture grade saline (0.9%, Sigma). The hippocampal region of the brain was isolated, and then dissociated into a viable single cell suspension using the Adult Mouse and Rat Brain Dissociation kit (Miltenyi) using a gentleMACS Octo Dissociator with Heaters automated tissue dissociation instrument. This process incorporates both enzymatic and mechanical digestion to remove viable cells from the extracellular matrix and removes debris through gradient centrifugation, taking a total of approximately 3 hours. Cells were cryopreserved using Recovery Cell Culture Freezing Medium (Thermo Fisher). Viability and cell concentrations were quantified upon thawing for single cell analysis using a Luna FL Automated Cell Counter (Logos) by co-quantification of fluorescence from acridine orange and propidium iodide dyes. We profiled eight total samples: four Pb treated and four control, with biological replicates from two male and two female animals in each treatment group. Cells were resuspended in a PBS + 0.02%BSA solution at a concentration of approximately 1000 cells/mL for single cell transcriptomic analysis processing.

### Single Cell RNA-sequencing

Single cells were processed for high throughput RNA sequencing using the Chromium (10× Genomics) instrument at the University of Michigan Advanced Genomics Core Facility, targeting approximately 3,000-5,000 cells per sample. Cells from the eight hippocampus samples were processed. The Chromium device partitions cells into individual oil droplets containing gel beads which are hybridized to oligonucleotides containing a partial Illumina sequencing primer, a unique molecular identifier (UMI), a poly-dT primer sequence for the capture of mRNA, and a cell specific 10× barcode. Chromium uses a pool of approximately 750,000 different 10× barcodes to index each cells transcriptome separately, with each gel bead having a single 10× barcode. Cells are lysed, and reverse transcription reactions conducted within the oil droplet result in cDNAs which incorporate a cell specific oligo barcode. The oil beads were broken and cDNAs from all cells are pooled together, the cDNA pool is amplified by qPCR, and Illumina P5 and P7 sequencing primers added during Sample Index pPCR. Prepared libraries for all 8 samples were pooled and sequenced on a NovaSeq 6000 (Illumina) using 2×96 paired end reads to capture sample index, cell barcode, and transcriptional information.

### Data Processing

Raw single cell RNA-sequencing data were processed using the CellRanger (10× Genomics) analysis pipeline. *mkfastq* was used to demultiplex the raw Illumina BCL files into fastq format. *count* was used to align the reads to the mouse reference genome (mm10), filter reads, count barcodes, and count UMIs. The output from *count* yields digital gene expression (DGE) matrices for each sample that contain the UMI counts per gene, per cell barcode. DGEs were loaded into the single cell transcriptomic data analysis suite Seurat (Butler et al. 2018), then normalized using the “LogNormalize” method in Seurat.

### Data Analysis

We tested for differences by treatment group in cell viability prior to processing on the 10X instrument using a t-test. For data quality control, we applied multiple filter criteria (**Supplemental Figure 1**). We excluded droplets with fewer than 1,000 expressed genes and we excluded genes that were measured in fewer than three cells, leaving a total of 5,258 cells and 17,143 genes expressed across the 8 samples. We calculated the proportion of mitochondrial genes expressed per cell. We tested for differences in quality control metrics by treatment using t-tests. Specifically, we tested for differences in the number of droplets sequenced, the proportion of cells passing quality control and following filtering, and the mean proportion of mitochondrial genes expressed.

Dimension reduction was performed on scaled DGEs using principal component analysis (PCA). For data visualization, t-Distributed Stochastic Neighbor Embedding (tSNE) (Maaten and Hinton 2008) was conducted on the top 12 principal components. The number of principal components was chosen heuristically based on an elbow plot (**Supplemental Figure 2**), where the variance explained by each principal component began to level out. Cell clusters were predicted based on the top 12 principal components using the *FindClusters* function, which first uses a k-nearest neighbors graph based on Euclidean distance in PCA space, followed by clustering by the Louvain algorithm. Key gene markers which define each cluster were calculated using the *FindAllMarkers* function, which uses the non-parametric Wilcoxon rank sum test to identify the top expression markers which are upregulated in each cell cluster. We assessed cluster identity using known markers of the various cell populations in the hippocampus as well as predicting each cluster’s identity based on the expression of cell type specific marker genes identified previously in single cell RNA-seq profiling of the mouse hippocampus (Zeisel et al. 2015).

We summarized the proportion of cells assigned to each cell cluster by animal. We collapsed cluster counts by our identified cell types, and for each cell type, we tested for differences in the proportion of cells by treatment (Pb vs. control) using beta regression. We examined differential expression analysis by Pb using edgeR. We used glmQLFit with a nested design to adjust for each individual mouse within the two treatment groups and also included the logit of each cell’s detection rate (proportion of non-zeros) as a recommended covariate (Soneson and Robinson 2018). To test for differential expression in Pb treatment, we used glmQLFTest with a contrast to compare the average effects of the mice in the Pb group to those in the control group. We then calculated false discovery rate (FDR) q-values to account for multiple comparisons. We tested for differential expression by Pb first with all cell clusters together. We also conducted a cell cluster specific analysis by subsetting cells from each individual cluster, then testing for differential expression by Pb. To investigate whether bulk analysis with all cells together reflects cell proportion changes, we tested the overlap of top differentially expressed genes and cluster markers.

We performed gene set enrichment analysis using the RNA-Enrich option of LRpath (Lee et al. 2016). P-values, effect estimates and log counts per million from the edgeR differential expression results were used as input for LRpath. For enrichment analysis we did directional tests in LRpath, querying the GO Biological Processes database. First, enrichment analysis was performed on differentially expressed genes in the analysis across all clusters. Cluster specific enrichment analysis was done on cluster specific differentially expressed genes.

## Results

### Sequencing of hippocampal cells

To identify effects of perinatal Pb exposure in the hippocampus on a cell-specific basis, we profiled eight 5-month old dissociated hippocampal samples (N=4 Pb exposed and N=4 controls; 50% male). The observed number of droplets sequenced ranged from 376,198 to 455,785 per animal (**Supplemental Table 1**). Following quality control filtering for number of genes expressed and percent mitochondrial genes expressed, we obtained results from 5,258 hippocampal cells across all animals. Per sample, the geometric mean percent mitochondrial genes expressed ranged from 3.0%−4.7% and was on average 0.79% lower with Pb exposure (p-value = 0.046) (**Supplemental Figure 3**). Given the proportion of mitochondrial genes were relatively low, we did not exclude cells on the basis of mitochondrial gene expression. Neither mean UMI nor mean genes expressed per cell differed significantly between control and exposed mice (UMI difference: 515.4, p-value = 0.09; genes per cell difference: 100.5, p-value = 0.15), although both trended higher in controls.

### Hippocampus cell cluster identification

Unsupervised cell clustering resulted in 12 cell clusters (**Figure 1A**). Clusters 0 through 5, and 7 through 9 had cells represented from each sample. Cluster 6 had no cells from two control samples and one Pb sample. Cluster 11 had no cells from one control sample. Cluster 10 cells were derived from one control sample (88 of 93 cells), and one Pb exposed sample (**Figure 1B**). Cell clusters ranged in size from 44 to 1329 cells (**Table 1**).

**Figure 1.**
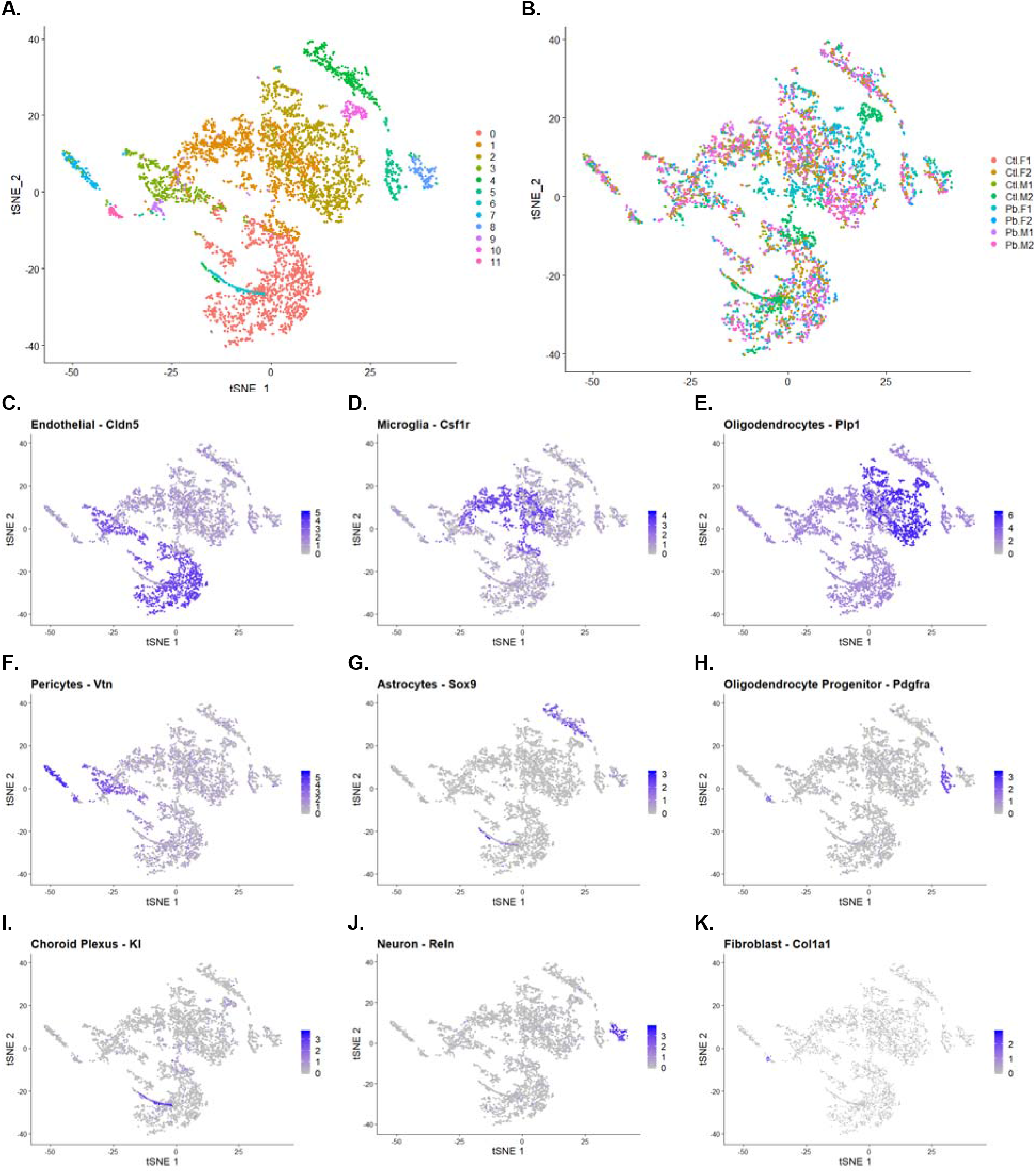
t-Distributed Stochastic Neighbor Embedding (tSNE) by cell cluster 0-11 (A) and sample (B). Samples are labeled as control samples (Ctl) and Pb-treated (Pb), the sex of the sample is specified (male (M) and female (F)) and biological replicates are indicated 1-2. Figures are painted by known markers for cell types: *Cldn5* for endothelial cells (C), *Csf1r* for microglial cells (D), *Plp1* for oligodendrocytes (E), *Vtn* for pericytes (F), *Sox9* for astrocytes (G), *Pdgfra* for oligodendrocyte precursors (H), *Kl* for choroid plexus (I), *Reln* for neurons (J), *Col1a1* for fibroblasts (K).

**Table 1.**
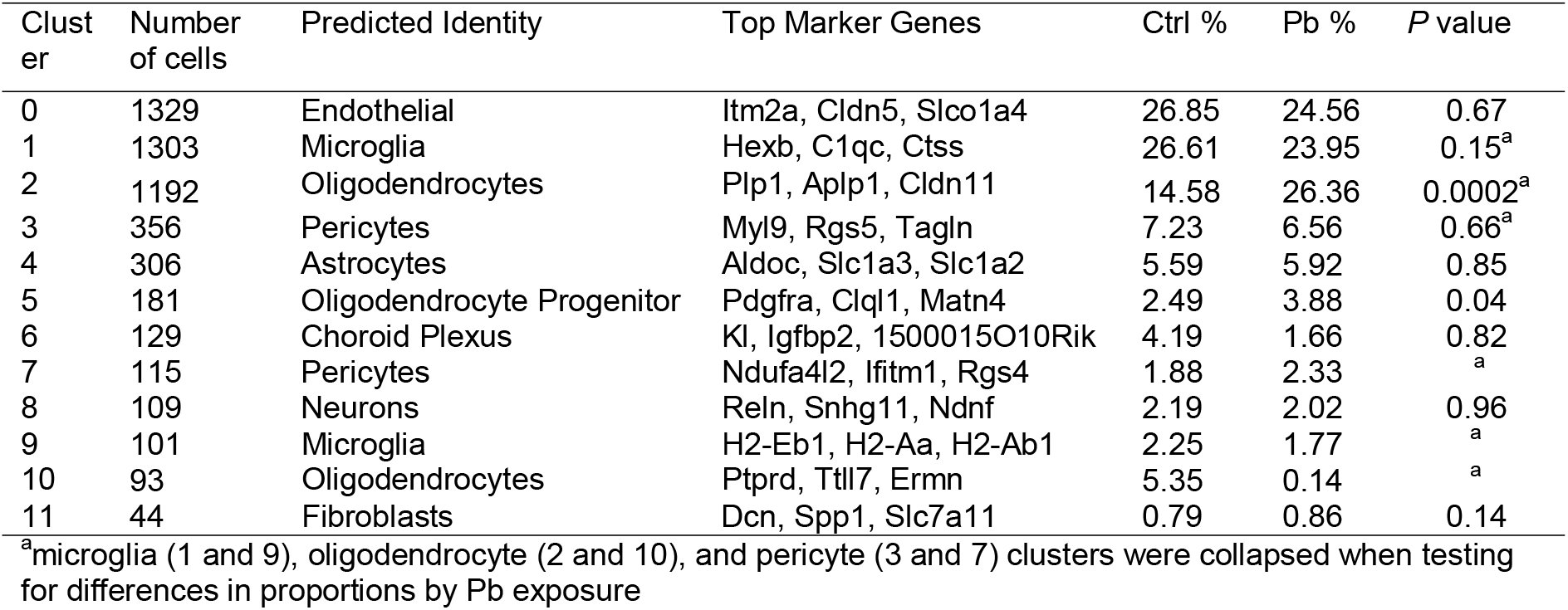
Number of cells observed per cluster and the predicted cell type identity of that cluster.

We identified likely cell types for each cluster using published marker genes (**Supplemental Table 2**). Endothelial cells were the most abundant cells, identified in cluster 0 by their expression of Claudin 5 (*Cldn5*) (**Figure 1C**). Microglial cells were the next most abundant cells, identified in clusters 1 and 9 by their expression of colony stimulating factor 1 receptor (*Csf1r*) (**Figure 1D**). Oligodendrocytes can then be observed in clusters 2 and 10 by expression of Proteolipid protein 1 (*Plp1*) (**Figure 1E**). Pericytes were identified in clusters 3 and 7 by their expression of vitronectin (*Vtn*) (**Figure 1F**). Astrocytes were identified in cluster 4 by their expression of SRY-box 9 (*Sox9*) (**Figure 1G**). Cluster 5 is likely oligodendrocyte progenitor by their expression of Platelet-derived growth factor receptor (*Pdgfra*) (**Figure 1H**). Klotho (*Kl*) expression marked choroid plexus cells in cluster 6 (**Figure 1I**). Neurons were identified in cluster 8 by their expression of Reelin (*Reln*) (**Figure 1J**). Lastly, fibroblasts were the least abundant cells, identified in cluster 11 by their expression of collagen type I alpha 1 (*Col1a1*) (**Figure 1K**).

The geometric mean number of genes expressed per cell varied by cell cluster (mean = 2537.2, range = 1658.3, 3619.4) (**Supplemental Figure 4**). Cell types with the highest number of genes expressed were choroid plexus, pericytes, oligodendrocyte progenitor cells, and neurons (clusters 3, 5, 6, 8). Cell types with the lowest number of expressed genes were microglia, astrocytes, and pericytes (clusters 1, 4, 7, 9).

### Overall differences in gene expression by Pb exposure

Across all cells (n_cells_ = 5,258) and all genes (n_genes_ = 17,143), we tested for differences in gene expression by Pb exposure (**Supplemental Table 4**). We observed 1,230 genes that differed by Pb exposure (q-value < 0.05), and of these, 5 had absolute log fold change greater than 0.5 (**Figure 2A**). Among these genes, only one (*Hapln2*) had higher expression with Pb exposure while four had lower expression with Pb exposure. Hemoglobin, beta adult s chain (*Hbb-bs*) was 0.74 log-fold lower expressed in Pb exposed animals (q-value=5.0×10^−72^). A predicted gene on chromosome 5, *Gm42726*, had 0.74 log-fold lower expression in Pb exposed animals than controls (q-value = 2.0×10^−40^) (**Figure 2B**). Kinesin family member 5A (*Kif5a*) was 0.65 log-fold lower expressed in Pb exposed animals than controls (q-value = 1.3×10^−33^) (**Figure 2C**). Hyaluronan and proteoglycan link protein 2 (*Hapln2*) was 0.56 log-fold higher expressed in Pb exposed animals than controls (q-value = 4.7×10^−17^) (**Figure 2D**). Predicted gene *Gm15013* on the X chromosome was 0.55 log-fold lower expressed in Pb exposed animals than controls (q-value = 7.8×10^−15^) (**Figure 2E**). These genes are not uniformly expressed by all cell types (**Supplemental Figure 5**).

**Figure 2.**
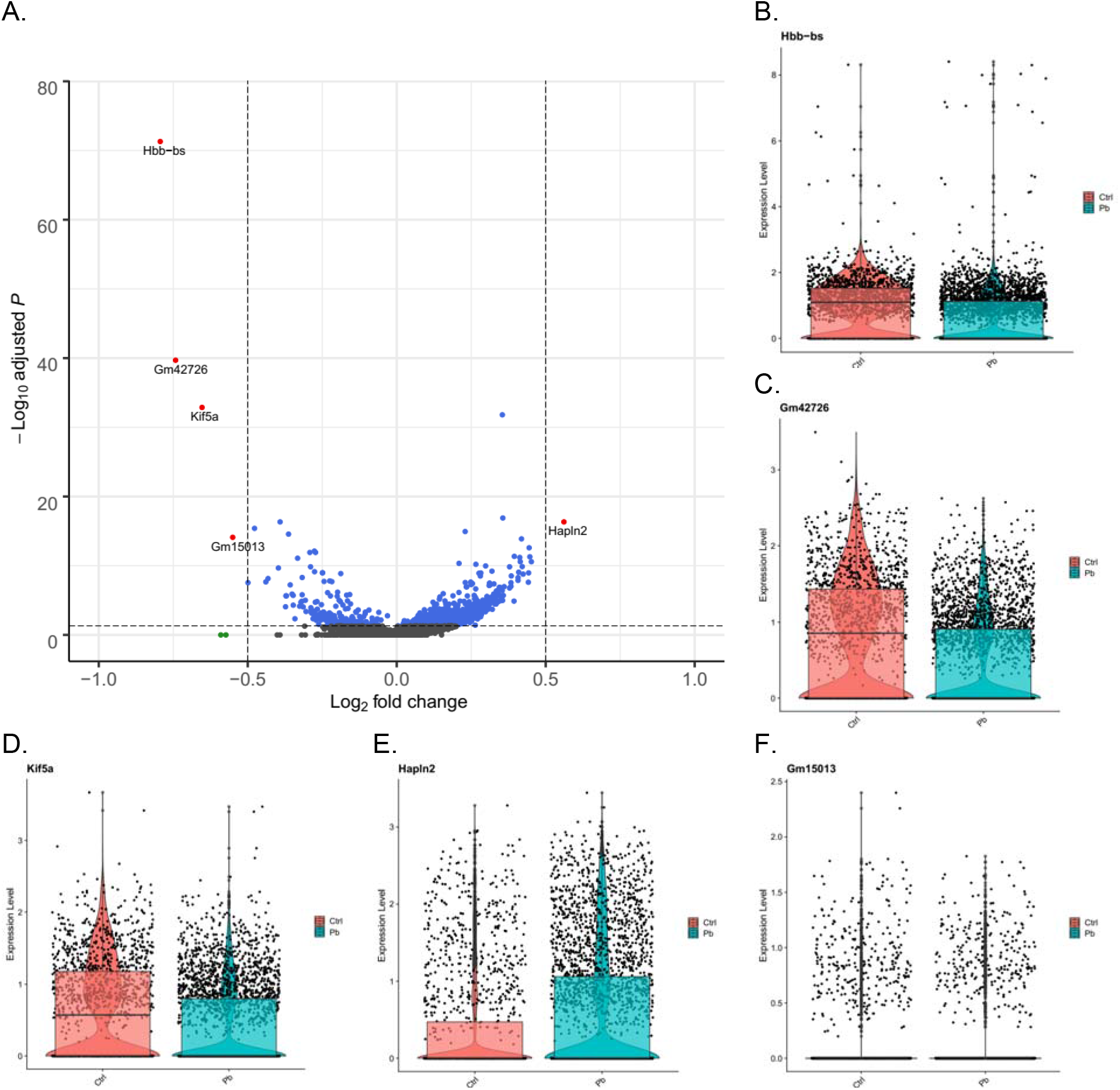
Across all cells measured, genes differentially expressed in the mouse hippocampus at 5-months of age following perinatal Pb exposure. Volcano plot of the log(fold change) difference in expression between Pb exposed and control versus the −log_10_(FDR q-value) for the test statistic. Each point represents a gene. Genes reaching q-value < 0.05 are blue, genes reaching log(fold change) > ± 0.5 are green, and genes reaching both thresholds are red (A). Violin plots showing per cell gene expression in Pb exposed animals versus controls for *Hbb-bs* (B) *Gm42726* (C), *Kif5a* (D), *Hapln2* (E), and *Gm15013* (F). FDR: False discovery rate.

To assess the contribution of cell cluster proportions to the overall Pb differences, we tested the overall Pb differentially expressed gene list for enrichment in cell cluster marker genes. Specifically, the top 10 markers for oligodendrocytes in cluster 2 (*Plp1*, *Aplp1*, *Cldn11*, *Cnp*, *Ptgds*, *Car2*, *Mag*, *Cryab*, *Mog*, *Mal*) and the top 10 markers for oligodendrocyte progenitor cells in cluster 5 (*Tnr*, *Lhfpl3*, *Cacng4*, *Cdo1*, *Gpr17*, *Neu4*, *3110035E14Rik*, *Opcml*, *Pcdh15*, *Vcan*) were among the Pb exposure bulk analysis 1,230 differentially expressed genes with q-value < 0.05 (Fisher p-value for overlap = 3.5×10^−12^). Several of the top 10 cluster markers for oligodendrocytes in cluster 10 (*Ptprd*, *Pcdh9*, *Kcna1*, *Edil3*, *Agpat4*, *Ttll7*, *Cntn2*) were also significantly associated with Pb in bulk analysis (Fisher p-value for overlap = 9.5×10^−7^).

### Pb exposure related changes in cell cluster proportions

We observed Pb exposure cells were non-randomly distributed across cell clusters (**Figure 3A**). We tested for differences in the proportions of cells annotated to cell types by treatment group (**Supplemental Figure 6**) with beta regression. The proportion of oligodendrocyte cells was higher on average with Pb exposure (p-value = 0.0002), and the proportion of oligodendrocyte progenitor cells was marginally higher with Pb exposure on average (p-value = 0.04) (**Table 1**).

**Figure 3.**
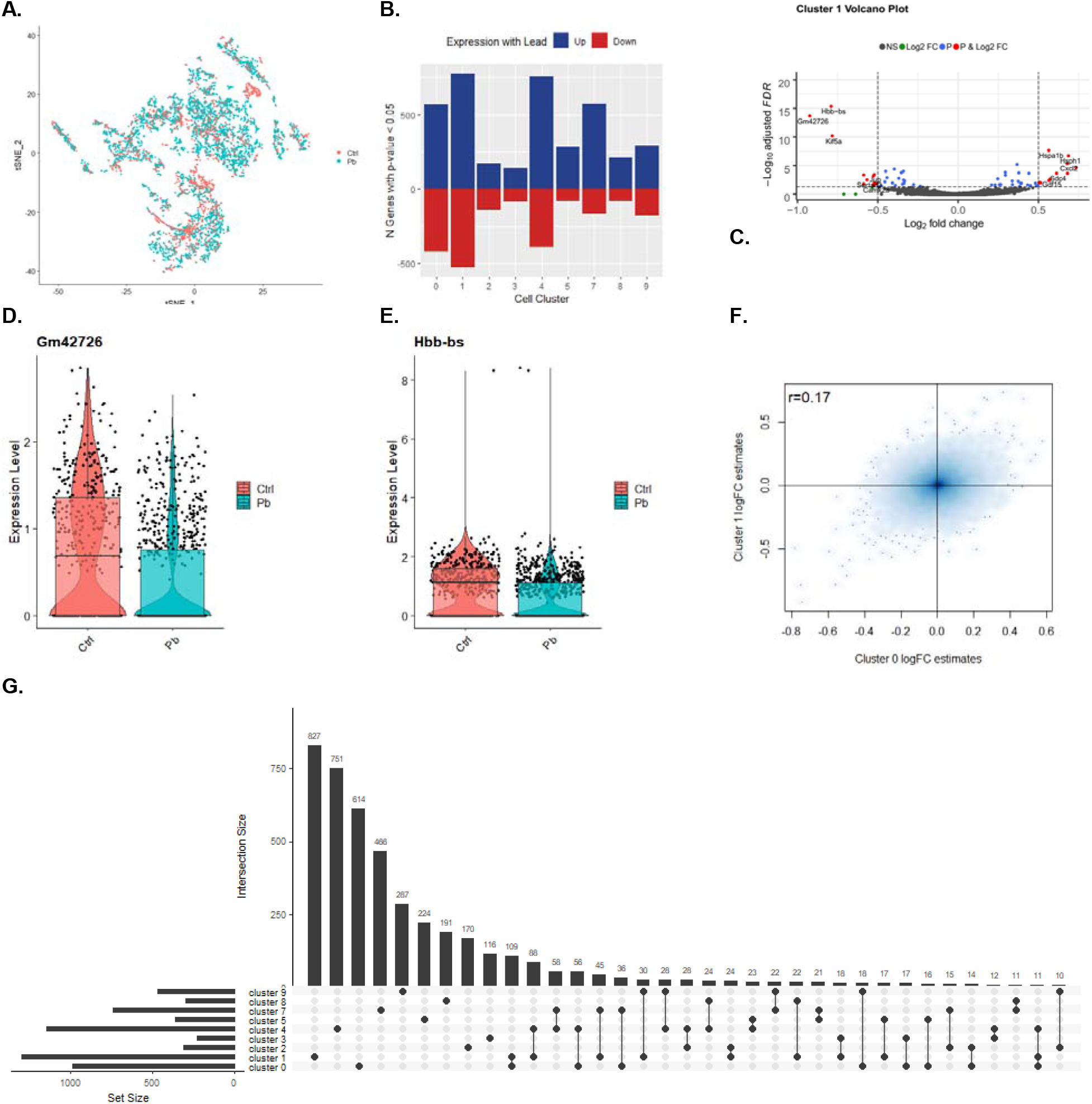
Cluster specific differences in gene expression by Pb exposure. tSNE by Pb exposure (A). Barchart of number of nominal (p-value < 0.05) differentially expressed genes by Pb exposure per cluster (B). Volcano plot of cell cluster 1 differential expression (C). Violin plots of gene expression levels in cluster 1 cells of Gm42726 (D). and Hbb-bs (E). Scatter plot of log(fold change) estimates in cluster 0 and cluster 1 (F). Upset plot of genes different/common across clusters with nominal p-value (<0.05), additional intersections with <10 genes not shown in plot (G).

### Pb exposure related differences within cell clusters

We tested for differences in gene expression by Pb exposure on a cluster specific basis (**Supplemental Table 4**). Due to zero cell counts for some samples in clusters 6, 10, and 11, we were unable to run tests in those clusters. Among the remaining clusters, we observed genes with differential expression by Pb exposure (q-value < 0.05 and absolute log(fold change) > 0.5) in four clusters: clusters 0, 1, 3, and 4 (**Figure 3B**). Microglia (cluster 1) had 22 genes differentially expressed with Pb (**Figure 3C**). The top genes were *Gm42726*, which had 0.92 lower log(fold change) expression by Pb (**Figure 3D**) and *Hbb-bs* which had 0.79 lower log(fold change) expression by Pb (**Figure 3E**). Endothelial cells (cluster 0) had 9 genes differentially expressed by Pb. Pericytes (cluster 3) had 1 such gene, and astrocytes (cluster 4) had 2 genes meeting those criteria (**Supplemental Figure 7**).

To assess the consistency of Pb’s effects across cell types, we tested the pairwise Spearman correlations of gene-specific effect estimates. Log(fold change) by Pb treatment were most correlated between endothelial (cluster 0) and microglia (cluster 1), with Spearman r=0.17 (**Figure 3F**). Pairwise log(fold change) estimate relationships among other clusters had correlations r<0.1, except for endothelial (cluster 0) versus pericytes (cluster 3) (r=0.16) and microglia (cluster 1) versus pericytes (cluster 3) (r=0.13) (**Supplemental Figure 8**). Most genes differentially expressed by Pb exposure within clusters were cluster specific, with comparatively few genes overlapping between clusters (**Figure 3G**). A total of 166 genes were nominally differentially expressed in three or more clusters. One gene, *Hbb-bs,* was nominal in all clusters, except in neurons (cluster 8).

### Gene set enrichment

Gene set enrichment by Pb exposure was performed on an overall and cluster specific basis. In our overall analysis (similar to a bulk RNA analysis), the top pathways in Pb exposure were upregulation of oligodendrocyte differentiation and ensheathment of neurons, and downregulation of lymphocyte activation (**Supplemental Table 5**).

The majority of genes implicated in cluster-specific pathway analyses were unique to a single cluster (**Figure 4A**). The top pathways in cluster specific analysis include enrichment in the following gene sets: downregulation of mRNA poly(A) tail shortening (endothelial cells in cluster 0), upregulation of protein folding (microglia in cluster 1, **Figure 4B**), upregulation of interleukin-12 biosynthetic process (oligodendrocytes in cluster 2, **Figure 4C**), down regulation of regulation of tyrosine phosphorylation of Stat5 protein (pericytes in cluster 3), upregulation of regulation of plasma lipoprotein particle levels pathway (astrocytes in cluster 4upregulation of positive regulation of stem cell differentiation pathway (oligodendrocyte progenitor cells in cluster 5), downregulation of double-strand break repair via nonhomologous end joining (pericytes in cluster 7), downregulation of phospholipase C-activating G-protein coupled receptor signaling pathway (neurons in cluster 8), and upregulation negative regulation of protein oligomerization (microglia in cluster 9, **Figure 4D**) (**Supplemental Table 6**).

**Figure 4.**
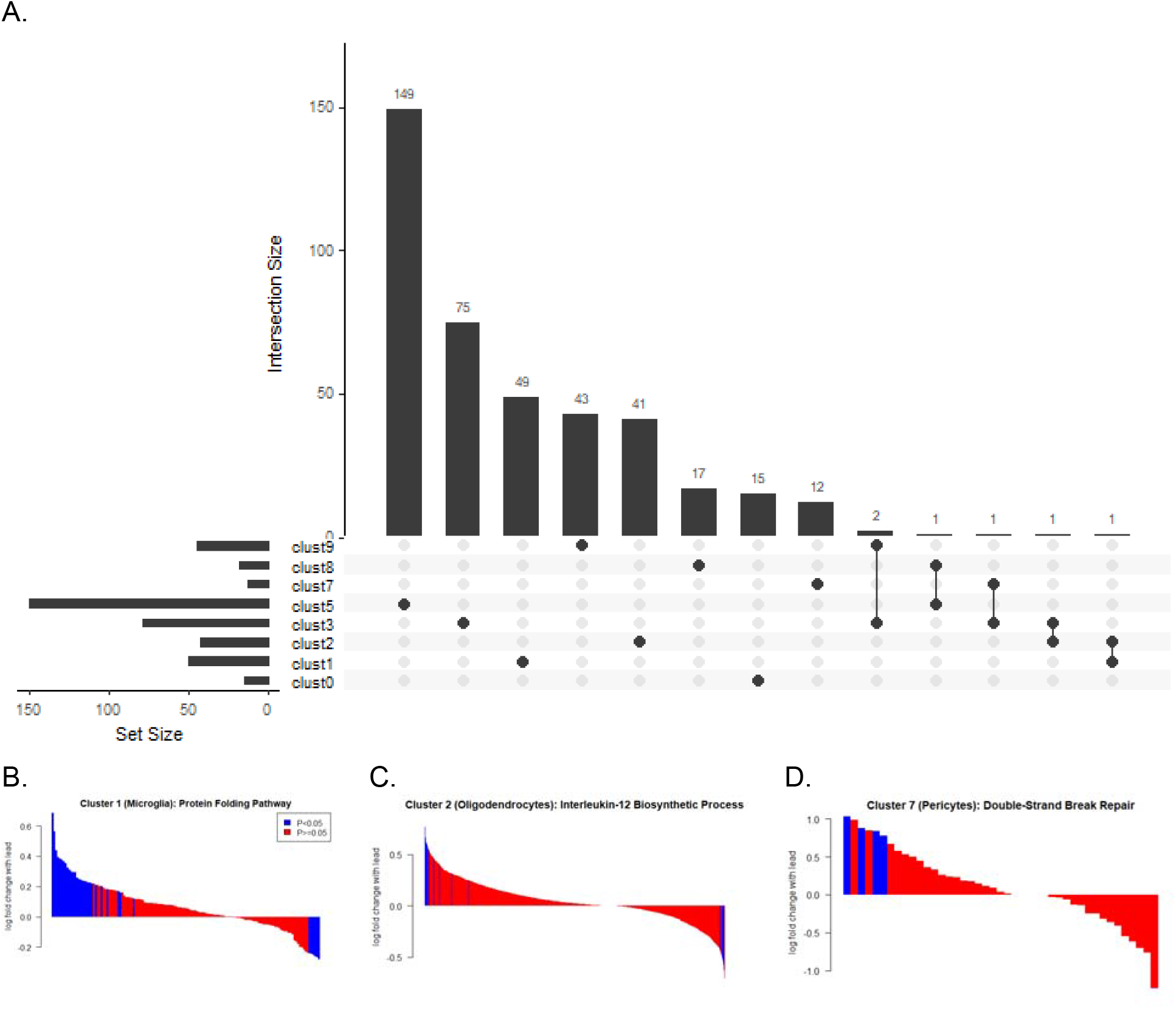
Gene set enrichment analysis on cluster-specific gene expression differences with Pb exposure in the hippocampus. In clusters 0, 1, 2, 3, 5, 7, 8, and 9, gene ontologies were identified with FDR<0.05 and these ontologies contained genes that were largely unique (A). In panels B-C, genes within a cluster are shown on the x-axis, ranked by their magnitude of association with Pb exposure and painted by the level of significance of their association with Pb exposure. In cluster 1, the protein folding pathway was upregulated (GO: 0006457) (B). In cluster 2, the interleukin-12 biosynthetic process pathway was upregulated (GO:0006950) (C). In cluster 7, double-strand break repair via nonhomologous end joining pathway was downregulated (GO:0006303) (D).

## Discussion

Toxicological and epidemiological evidence shows that early life exposure to Pb has profound impacts on the developing brain, with no safe level of Pb exposure identified. The mechanisms and effects of Pb’s effects in the brain, however, remain incompletely understood. Here, we used single cell RNA-sequencing to compare gene expression patterns in adult mouse hippocampus following perinatal Pb exposure. These methods allow for unparalleled resolution of cell type specific effects of Pb exposure. Through unbiased clustering and marker gene analyses, we identified and separated multiple known cell types in the brain, including oligodendrocytes, endothelial cells, microglia, neurons, pericytes, astrocytes, and fibroblasts. We found that *in utero* Pb exposure shifts the proportions of these cells, and also causes cell type specific transcriptomic alterations which persist into adulthood. Many of these changes were in genes and pathways associated with neurodegenerative diseases, as well as in some novel genes and pathways not previously associated with Pb exposure. Overall, these findings highlight the power of single cell analyses for the assessment of the cell type specific effects of toxicant exposure and provide new mechanistic insights into the lifelong effects of developmental Pb exposure.

We identified significant shifts in adult hippocampal cell type proportions following perinatal Pb exposure. Specifically, Pb exposed animals had a significantly higher proportion of oligodendrocytes relative to control animals. Additionally, Pb exposed animals also had a higher proportion of oligodendrocyte progenitor cells. Our findings of oligodendrocytes being a particularly Pb-susceptible cell population corroborates others’ prior work in this area. *In vitro* treatment of oligodendrocyte progenitor cells with Pb caused decreased differentiation capacity, decreased myelin basic protein (MBP) expression, and decreased cell branching (Ma et al. 2015). Wistar rats exposed to 1% Pb in their drinking water for 3 to 6 months had significant hypomyelination and demyelination in their nerve fibers (Coria et al. 1984), reflecting aberrant oligodendrocyte function. *In vitro* studies show that oligodendrocyte progenitor cells are more susceptible to the effects of Pb than differentiated oligodendrocytes (Deng et al. 2001), providing further mechanistic evidence that link developmental Pb exposures to altered neurological function. In agreement with these previous studies, we also identified an increase in oligodendrocyte progenitor cells in Pb exposed animals. Additional time course studies could reveal whether these changes were due to an expansion of oligodendrocytes early in life or due to a depletion of other cell populations in the brain. Further research is necessary to determine whether early life exposure to Pb causes an expansion of oligodendrocytes as a compensatory mechanism for decreased myelination capacity, or if these increased proportions of oligodendrocytes come due to a relative depletion of other cell types in the hippocampus.

Of relevance to interpreting results from bulk tissue molecular profiling studies in toxicology, we also simulated the standard approach in the field for differential expression analysis by comparing gene expression across bulk tissues in treated and control animals. In these analyses, we identified overall gene expression differences with Pb exposure that were driven by differences in cell proportions, specifically greater numbers of oligodendrocytes with Pb exposure. Indeed, the top cell type markers for multiple clusters, including oligodendrocytes and oligodendrocyte progenitor cells predominated in the list of differentially expressed genes. These results highlight the importance of considering cell type heterogeneity and alterations in cell type proportions as major drivers of differences in bulk tissues molecular profiles between treated and control animals. Moreover, bulk tissue studies can be prone to contamination from surrounding tissue. In only one of our samples, we identified a gene expression signature of the choroid plexus, a brain region adjacent to the hippocampus. We suspect that these additional cells were captured during the dissection process of this sample. Previous studies of differences in hippocampal gene expression by treatment have identified known markers of the choroid plexus, including *Ttr*, *Igfbp2,* and *Kl* as the top differentially expressed genes (Sarvari et al. 2015). Our results suggest that these findings could actually reflect differential hippocampus contamination with choroid plexus tissue in the experimental groups. The precision to be able to detect and isolate this contamination is a significant advantage of single cell RNA-seq analyses in toxicology studies.

Many of our findings of genes and pathways differentially expressed with *in utero* Pb exposure align with recent literature linking altered gene expression, genetic variants, or known biological mechanisms associated with developing neurodegenerative diseases. For example, we identified that protein folding pathways were significantly downregulated in microglial cells with Pb treatment. Dysregulation of protein folding is hypothesized to be a major underlying etiologic factor for the development of Alzheimer’s disease (Selkoe 2003; Stocker et al. 2019). One of our top downregulated genes overall, and within individual clusters, was *Hbb-bs.* A genetic mouse model for Alzheimer’s disease, the Tg2576 mouse has lower levels of *Hbb-bs* in their hippocampus relative to wild type mice (Wu et al. 2019), suggesting that this gene may play a role in Alzheimer’s pathology in the hippocampus. We also identified significant downregulation of *Kif5a,* a kinesin gene involved in organelle transport. KIF5A was recently found to be downregulated in post-mortem temporal lobe samples from Alzheimer’s patients (Wang et al. 2019). This downregulation of KIF5A impaired the axonal transport of mitochondria and mitochondrial trafficking in the neuron. *KIF5A* was also identified in a recent genome-wide association study of amyotrophic lateral sclerosis (AML) (Nicolas et al. 2018). The finding of KIF5A alterations in both AML and Alzheimer’s, and in our perinatally Pb exposed mice point to this gene as a potential mediating factor linking early life Pb exposures to the development of these neurodegenerative diseases later in life. Alzheimer’s like pathology has been identified in aged non-human primates which were exposed to Pb starting at birth until 400 days of age (Wu et al. 2008). In addition, we observed upregulated plasma lipoprotein pathways in astroctyes. Lipoprotein levels and polymorphisms in Apolipoprotein E are hallmarks of Alzheimer’s disease risk (Liu et al. 2013). DNA damage is another factor in Alzheimer’s disease, where high levels of double strand breaks are observed in Alzheimer’s patients (Adamec et al.; Shanbhag et al.). This is consistent with elevated double strand break repair pathways in pericytes in our study. Surprisingly, we were able to these detect Alzheimer’s and AML-associated molecular alterations at only five months of age in mice. Our results suggest that molecular alterations due to the effects of Pb exposure persist into early adulthood and may precede histopathological alterations later in life. This work identifies an intervention point which may alter neurodegenerative disease trajectories for those exposed to neurotoxicants early in life to reduce risk of later life disease outcomes.

Our results should be interpreted in light of a number of study limitations. Due to the nature of single cell transcriptomic profiling experimental platforms, we were only able to assess effects in a small number of samples. Single cell transcriptomic studies are also conducted on dissociated tissue samples, which lose the spatial architecture of the tissue. Moreover, droplet based platforms require viable cells and have differential capture rates based on cellular characteristics, such as shape and size, or genes expressed (Ye et al. 2019). For example, here we captured an overabundance of glial cells relative to neurons. Future single cell studies of the neuron specific effects of toxicant exposure could first capture neurons or neuronal nuclei using a fluorescence activated or magnetic bead-based sorting method to ensure a population of enriched cells for downstream analysis. Emerging literature also identifies background contamination of highly expressed transcripts (a so-called “soup” effect) that is detectable across all droplets (Young and Behjati 2018), likely due to dying cells releasing RNA into the buffer solution being run through the Chromium instrument, which are then detectable in all droplets containing that buffer. Ongoing improvements in experimental technology and bioinformatic analysis methods will likely dramatically increase the utility of single cell transcriptomics for toxicology studies. Future studies could also integrate orthogonal validation techniques, such as immunohistochemistry analysis or RNA *in situ* hybridization approaches in intact tissues. These analyses were not available for the mice in this study, which were part of the TaRGET II consortium. All tissues were earmarked for high throughput ‘omics analyses, rather than pathologic assessment. Ongoing analyses by our group, including whole transcriptome and methylome profiling of other brain tissues, are current underway for integration with these single cell data.

Our study also has a number of strengths. To the best of our knowledge, this is the first single cell analysis of the effects of a neurotoxicant on the brain. This approach allowed us to simultaneously assess the effects of Pb on the cell type proportions of the developing brain as well as identify cell type specific effects. Deconvoluting the effects of a toxicant is essential for identifying mechanisms of action and biomarkers of exposure, for example, identifying whether an exposure during development alters the cell type proportions in the adult organ or alters gene expression patterns of individual cell types in a tissue. Our study also used an established exposure paradigm to understand the effects of perinatal toxicant exposure conducted in the context of a large national consortium. Overall, these results inform our understanding of the molecular effects of early life Pb exposure as well as the interpretation of data from bulk tissue RNA-seq data in toxicology experiments.

Pb exposure remains a persistent public health challenge in the US and worldwide. Our results here use single cell transcriptomic profiling to help to elucidate the molecular effects of early life exposure to Pb. We identify that glial cell populations, particularly cellular proportions of oligodendrocytes and gene expression in microglia, appear to be the most susceptible to the effects of Pb, and that these effects persist into adulthood. Neurons were underrepresented in our sample. We also identified a number of important genes and pathways simultaneously associated with neurodegenerative diseases and Pb exposure. Future work should characterize the mechanism by which these alterations lead to frank pathology in the brain through aging, whether through epigenetic alterations, oxidative stress, or persistently activated inflammatory pathways. Cutting edge techniques, including single cell RNA-seq and spatial transcriptomic profiling, will continue to provide new insights to these biological processes at unprecedented resolution.

## Acronyms

FDR: false discovery rate
UMI: unique molecular identifier
DGE: digital gene expression
FC: fold change
GO: gene ontology
Pb: lead
PCA: principal component analysis
tSNE: t-Distributed Stochastic Neighbor Embedding
(TaRGET) II consortium: Toxicant Exposures and Responses by Genomic and Epigenomic Regulators of Transcription

## Acknowledgements

The mouse exposure study was supported by the NIEHS TaRGET II Consortium award to the University of Michigan (U01 ES026697). The sequencing experiment was supported by the NIEHS Michigan Center on Lifestage Environmental Exposures and Disease (M-LEEaD; P30 ES017885). Mr. Dou and Dr. Bakulski were supported by NIH grants (R01 ES025531, R01 ES025574, R01 AG055406, R01 MD013299). Dr. Colacino was supported by NIEHS (R01 ES028802). Dr. Perera was supported by NIEHS (T32 ES007062) and Dr. Neier was supported NIEHS (T32 ES007062) and by NICHD (T32 HD079342).

