## Supplementary Material for "Single cell analysis of the effects of developmental lead (Pb) exposure on the hippocampus"

**Title**

**Supplemental material**

27,998 overall genes measured

17,143 genes measured

Exclude 10,855 genes with < 3 cells expressed overall

3,353,993 overall droplets measured

(mean 419,249 per animal)

5258 cells measured

(mean 657 per animal)

Exclude 3,348,735 droplets with < 1000 genes expressed

(mean 418,592 per animal)

**A.	B.**

**Supplemental Figure 1**. Flow charts depicting inclusions and exclusions of droplets (**A**) and genes (**B**).

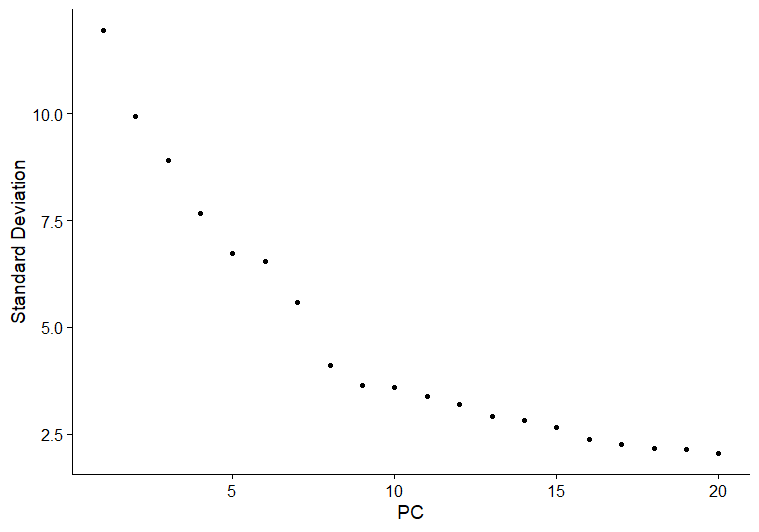

**Supplemental Figure 2**. Elbow plot

**Supplementary Table 1**. Single cell RNA sequencing data quality metrics by sample. Last three columns follow cell filtering.

|  | |  |  |  |  |  |  |  | Post-filtering, Per cell metrics | | |
| --- | --- | --- | --- | --- | --- | --- | --- | --- | --- | --- | --- |
| Sample | |  | Treatment | Sex | Initial Cell Viability | Droplets Sequenced N | Droplets Filtered N (%) | Cells Pass QC N | Number of Genes Expressed, Median (IQR) | Number of UMI, Median (IQR) | Percent Mitochondrial Genes Expressed, Median (IQR) |
| t103c | Pb.F1 | | Pb | F | 76.4% | 449009 | 447818 (99.73) | 1191 | 2401 (1189.5) | 6219 (4662) | 3.3 (1.7) |
| t107d | Pb.M1 | | Pb | M | 75.3% | 455785 | 454691 (99.76) | 1094 | 2466.5 (1189) | 6172.5 (4738.25) | 2.9 (1.7) |
| t109d | Ctl.M1 | | Ctl | M | 76.6% | 376198 | 375829 (99.9) | 369 | 2331 (1206) | 5855 (4379) | 4 (1.3) |
| t112c | Pb.F2 | | Pb | F | 77.7% | 438976 | 438502 (99.89) | 474 | 2280 (1230) | 5655.5 (4742.5) | 3.4 (1.6) |
| t113c | Ctl.F1 | | Ctl | F | 82.3% | 388256 | 388042 (99.94) | 214 | 2407 (1346) | 6387.5 (5703) | 4.5 (1.4) |
| t116d | Pb.M2 | | Pb | M | 72.5% | 416591 | 415738 (99.8) | 853 | 2292 (1088) | 5526 (4170) | 3.3 (1.6) |
| t118d | Ctl.M2 | | Ctl | M | 82.2% | 432571 | 432083 (99.89) | 488 | 2536 (1407.5) | 6536.5 (5369.5) | 4 (2.7) |
| t119g | Ctl.F2 | | Ctl | F | 78.5% | 396607 | 396032 (99.86) | 575 | 2500 (1353) | 6462 (4886.5) | 3.3 (1.2) |

**Supplemental Figure 3.** Violin plots by sample where each dot represents a cell. Post-filtering number of genes expressed per cell (A), number of unique molecular identifiers per cell (B), percent mitochondrial genes expressed per cell (C).

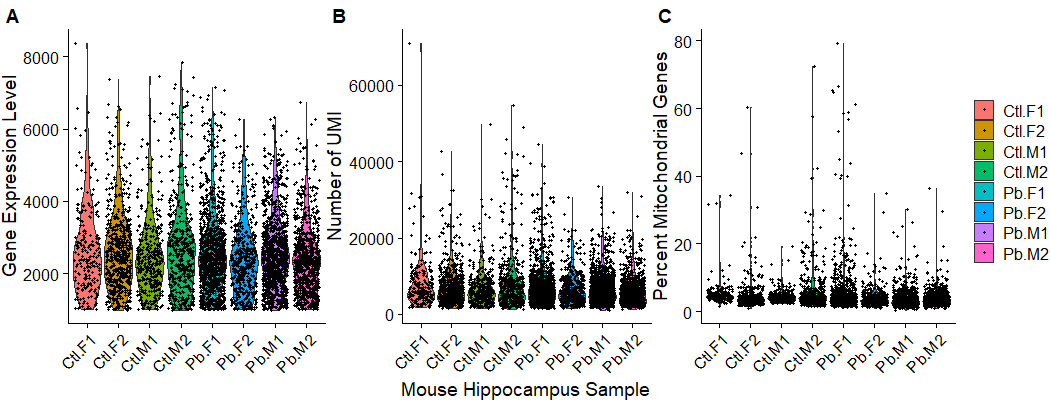

**Supplemental Table 2.** Marker genes by cluster.

Excel file, tab for each cluster showing genes with padj<0.05: Stable2_Cluster_markers_padj05.xlsx

**Supplemental Figure 4.** Violin plots by cell clusters. Post-filtering number of genes expressed per cell (A), number of unique molecular identifiers per cell (B), percent mitochondrial genes expressed per cell (C).

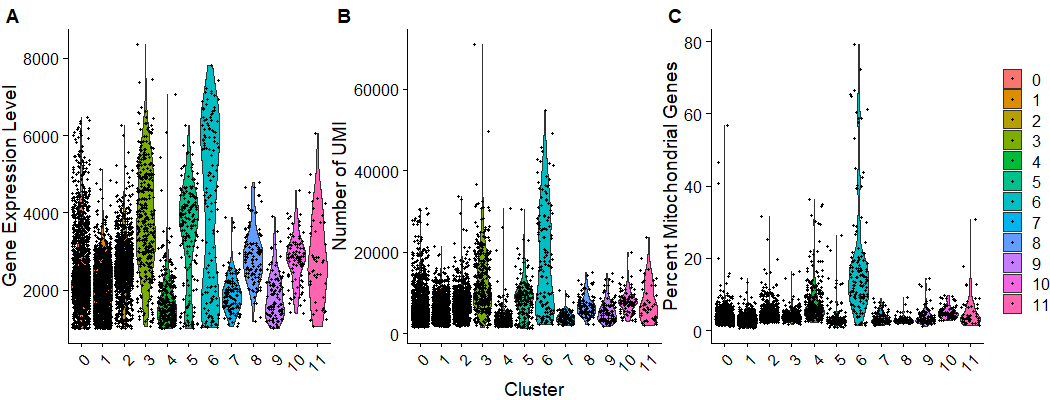

**Supplemental Table 3.** Full differential gene expression across all clusters by lead exposure.

CSV table: STable3_Pb_DE_genes_across_all_clusters.csv

**Supplemental Figure 5.** TSNE painted by top differential genes expressed across all clusters by lead exposure. Multi-panel.

| 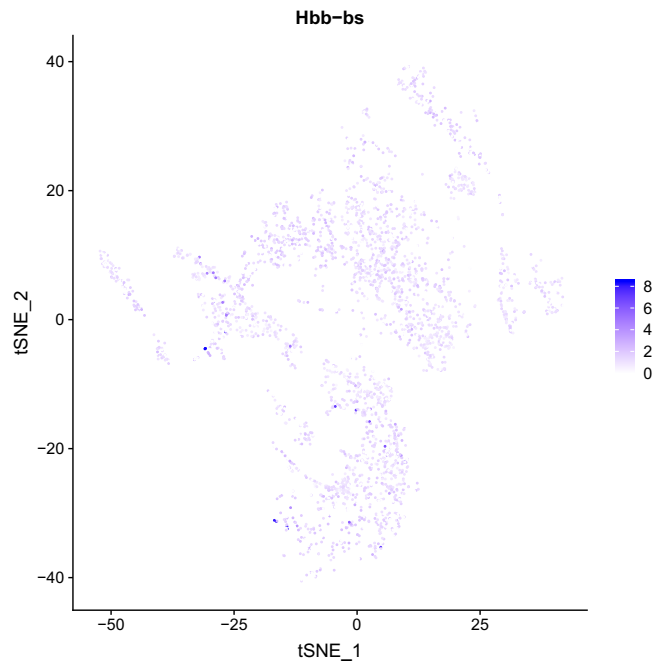 | 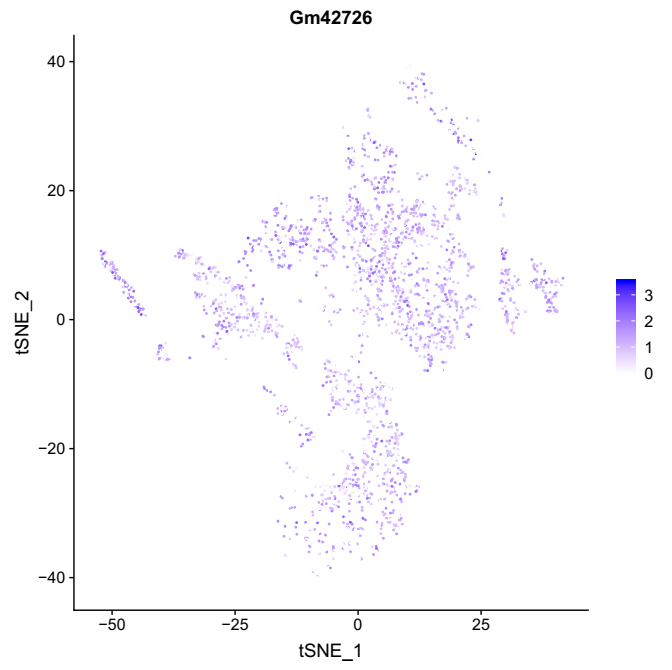 | 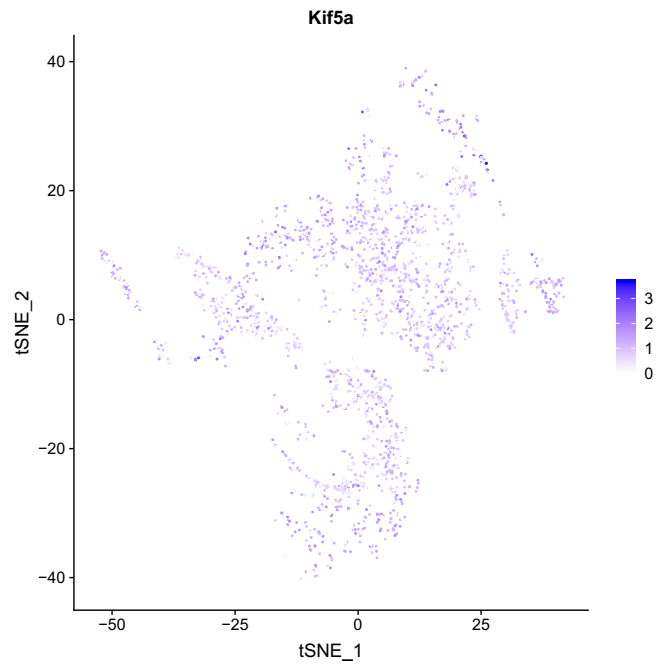 |
| --- | --- | --- |
| 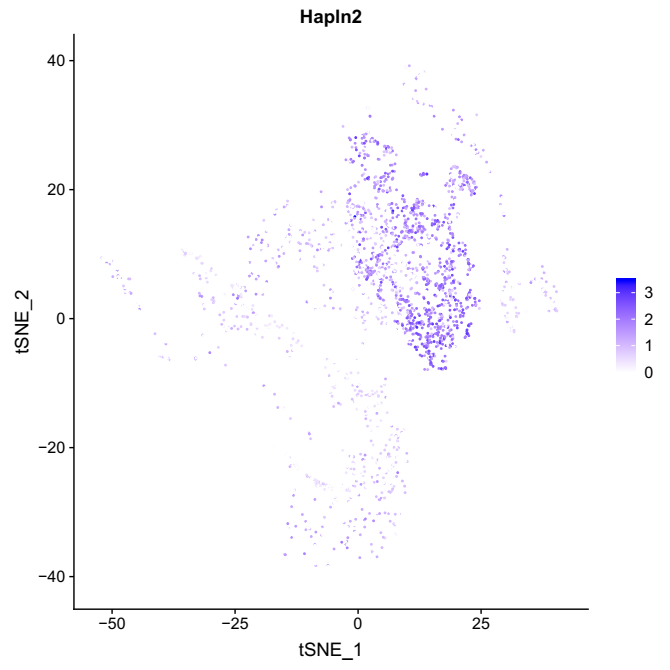 | 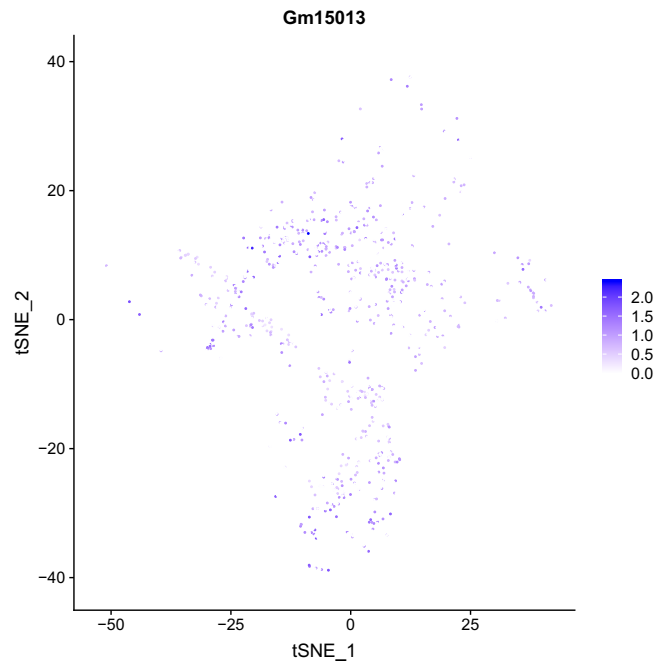 | 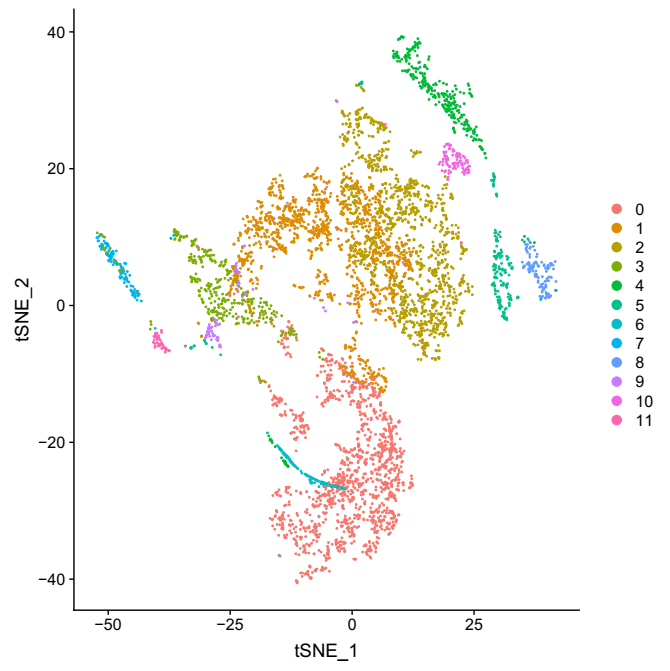 |

**Supplemental Tables 4.** Differential gene expression by cluster by lead exposure.

Excel file for all genes all clusters: STable4_Pb_DE_genes_cluster_specific.xlsx

**Supplemental Figure 6.** Stacked bar chart of proportions of presumed cell types by mouse. Clusters that were identified as the same cell type were collapsed: microglia (clusters 1 and 9), oligodendrocytes (clusters 2 and 10), and pericytes (clusters 3 and 7).

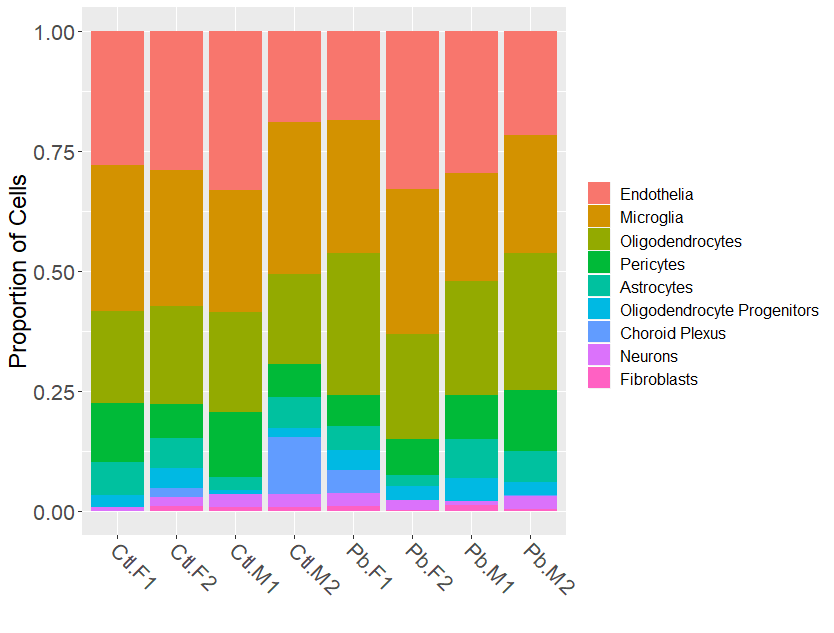

**Supplemental Figure 7**. Multi-panel plot of volcano plots for each cell cluster. Cutoff for log fold-change is 0.05. P-value cutoffs were set at maximum p-value if no genes had FDR<0.05, or at largest p-value that had FDR<0.05. Points are gray if genes do not meet log fold-change or p-value cutoff, green if they only meet p-value cutoff, blue if genes only meet log fold-change cutoff, and red if genes meet both log fold-change and p-value cutoff.

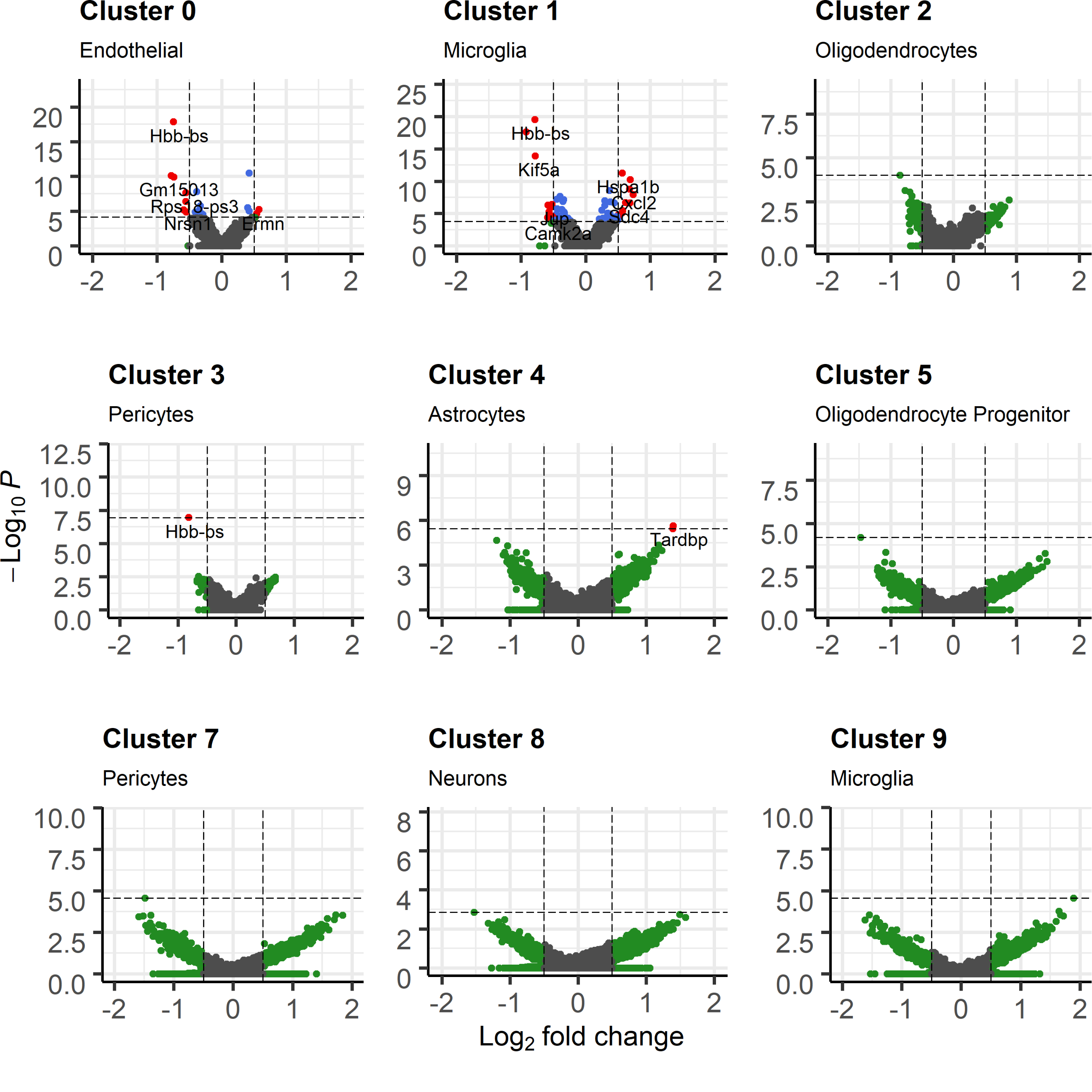

**Supplemental Figure 8.** Multi-panel scatter plot of fold change in one cluster versus another. Upper right panel shows the Spearman correlation coefficients. Lower left shows the Spearman correlation p-values.

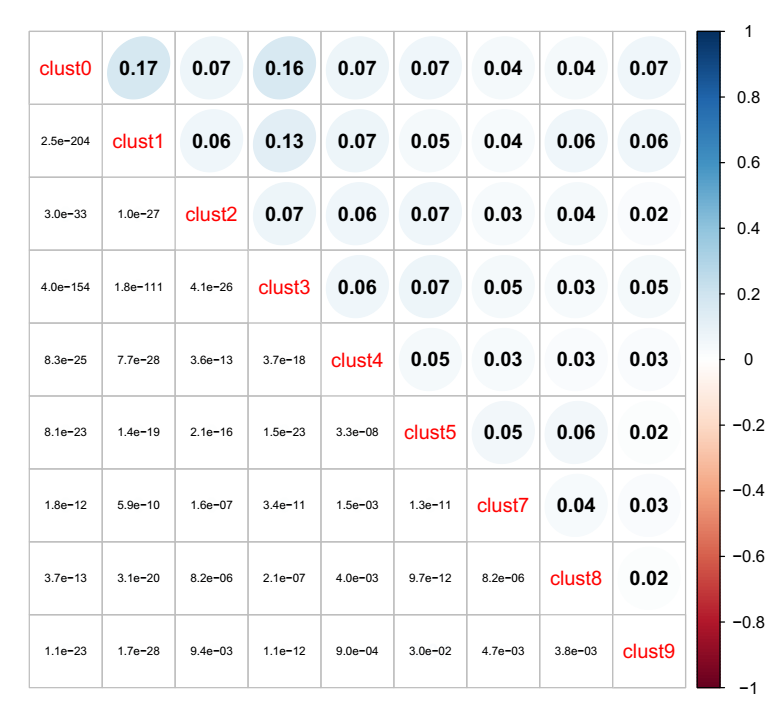

**Supplemental Table 5.** Gene ontology for genes nominally associated with lead exposure in bulk analysis of all clusters together.

File: STable5_LR_path_overall.xlsx

**Supplemental Table 6.** Gene ontology for genes nominally associated with lead exposure in cluster specific analysis.

File: STable6_LR_path_clust_specific.xlsx
